# Natural selection influenced the genetic architecture of brain structure, behavioral and neuropsychiatric traits

**DOI:** 10.1101/2020.02.26.966531

**Authors:** Frank R Wendt, Gita A Pathak, Cassie Overstreet, Daniel S Tylee, Joel Gelernter, Elizabeth G Atkinson, Renato Polimanti

## Abstract

Natural selection has shaped the phenotypic characteristics of human populations. Genome-wide association studies (GWAS) have elucidated contributions of thousands of common variants with small effects on an individual’s predisposition to complex traits (polygenicity), as well as wide-spread sharing of risk alleles across traits in the human phenome (pleiotropy). It remains unclear how the pervasive effects of natural selection influence polygenicity in brain-related traits. We investigate these effects by annotating the genome with measures of background (BGS) and positive selection, indications of Neanderthal introgression, measures of functional significance including loss-of-function (LoF) intolerant and genic regions, and genotype networks in 75 brain-related traits. Evidence of natural selection was determined using binary annotations of top 2%, 1%, and 0.5% of selection scores genome-wide. We detected enrichment (*q*<0.05) of SNP-heritability at loci with elevated BGS (7 phenotypes) and in genic (34 phenotypes) and LoF-intolerant regions (67 phenotypes). BGS (top 2%) significantly predicted effect size variance for trait-associated loci (σ^2^ parameter) in 75 brain-related traits (β=4.39×10^−5^, *p*=1.43×10^−5^, model *r*^*2*^=0.548). By including the number of DSM-5 diagnostic combinations per psychiatric disorder, we substantially improved model fit (σ^2^ ~ B_Top2%_ × Genic × diagnostic combinations; model *r*_*2*_=0.661). We show that GWAS with larger variance in risk locus effect sizes are collectively predicted by the effects of loci under strong BGS and in regulatory regions of the genome. We further show that diagnostic complexity exacerbates this relationship and perhaps dampens the ability to detect psychiatric risk loci.

## Introduction

Genome-wide association studies (GWAS) have identified numerous common genetic variants underlying thousands of human health and disease phenotypes.^1^ Psychiatric, neurological, and mental health related disorders and phenotypes (*e.g.*, personality and social science outcomes) have been investigated by large-scale GWAS, revealing substantial polygenicity, with small effects of loci across the genome contributing additively to a phenotype.^2^ Loci conferring risk or protection also have been shown to have a high degree of pleiotropy, with alleles associated with multiple traits through similar mechanisms.^3; 4^ For example, years of education has been negatively genetically correlated with *smoking behavior*,^3^ *body mass index*,^3^ *coronary artery disease*,^3^ *attention deficit hyperactivity disorder* (*ADHD*),^5^ *anxiety disorders*,^5^ *major depressive disorder*,^5^ and *stroke*,^5^ but is positively genetically correlated with *high density lipoproteins*,^3^ *anorexia nervosa*,^5^ *autism spectrum disorder*,^5^ *bipolar disorder*,^5^ *obsessive compulsive disorder*,^5^ and *schizophrenia*.^5^ The complex relationship between the shared polygenic architecture of presumably beneficial (*e.g., high education*) and debilitating (*e.g., schizophrenia*) traits intensifies a longstanding discussion about the origin and persistence of genetic risk for psychiatric and neurological disorders in the general population. One example of this puzzle is demonstrated with genetic risk for *schizophrenia*, a disorder observed in approximately 1% of the general population.^6^ *Schizophrenia* is often genetically and phenotypically associated with reduced fitness and fecundity.^7; 8^ Several hypotheses around this observation suggest that *schizophrenia* risk loci (1) confer advantage to those unaffected by the disorder or (2) are in strong linkage disequilibrium with positively selected alleles, resulting in an evolutionary trade-off between *schizophrenia* risk and more beneficial traits (*e.g., high intelligence*).^7; 9–11^

Background selection (BGS), the selective removal of alleles across the genome that confer deleterious effects, has been widely detected in the loci associated with complex traits, including those related to mental health and behavior.^12–16^ The effects of BGS are visible in the contribution of common and rare complex trait risk loci to overall trait heritability estimates: complex trait heritability is generally distributed evenly in common variants across the genome but is nonuniformly distributed across rare variants.^13^ These recent findings suggest that BGS has directly contributed to the degree of polygenicity of complex phenotypes and this relationship appears to be particularly strong in brain-related traits.^13; 16^ Because of the extremely complex genetic architecture of multifactorial brain-related phenotypes, the sample sizes needed to maximize explainable trait heritability exceeds several million.^2^ Even estimating these required sample sizes is further complicated by the effects of phenotype heterogeneity, which serves to reduce power for the detection of loci in GWAS, and is a systematic feature of diagnosed mental health conditions such as psychiatric disorders.^17; 18^

Many phenotypes of interest for large-scale GWAS have specific diagnostic criteria; for psychiatric disorders it is standard practice to refer to the Diagnostic and Statistical Manual of Mental Disorders, 5^th^ Edition (DSM-5). In practice, multi-item symptom check lists based on diagnostic criteria can be employed to classify an individual as a case *versus* a control. In performing this type of phenotype reduction, underlying differences in symptomology have the potential to produce large degrees of heterogeneity among the individuals being tested. For example, there are 636,120 possible combinations of symptoms which may lead to a *posttraumatic stress disorder* (*PTSD*) diagnosis by DSM-5 guidelines.^19^ Similar observations have been made with *ADHD*^20^ and *substance use disorders*.^21; 22^ While the symptoms used to define a specific diagnosis are generally highly correlated, these differences may still affect our ability to detect genetic signals associated with phenotype defined by diagnostic criteria.

Since gene discovery can provide information about the disease biology and identify potential treatment targets, it is imperative to study how evolution and diagnostic heterogeneity independently and interactively contribute to the polygenicity of traits related to mental health and disease. The goal of the present study was to investigate how natural selection influences polygenicity across 75 brain-related traits. With respect to psychiatric disorders, we investigated how the addition of heterogeneous trait definitions further contribute to our ability to detect genomic risk loci in psychiatric disorders. We find that both BGS and clinical heterogeneity increase the variance of effect size for risk loci of complex traits, including diagnosed psychiatric disorders, personality traits, internalizing and externalizing behaviors, social science trait outcomes, and brain imaging phenotypes.

## Subjects and Methods

### Datasets

GWAS summary association data were accessed from the Psychiatric Genomics Consortium (PGC), Social Science Genetic Association Consortium (SSGAC), UK Biobank (UKB), and UKB Brain Imaging Genetics (BIG) Consortium. All datasets were formatted as standard input for Linkage Disequilibrium Score Regression (LDSC). Observed-scale SNP-based heritability (*h*^*2*^) was calculated for all traits. For case/control phenotypes, *h*^*2*^ was calculated using effective sample sizes and converted to liability scale using population and sample prevalences reported in the respective publications (Table 1). The 1000 Genomes Project Europeans^23^ were used as the LD reference panel. Traits were selected for partitioning (see *Partitioned Heritability*) based on *h*^*2*^ z-score ≥ 7 (Table S1) as previously recommended.^24^

**Table 1.** Functional annotation enrichments. Three phenotypes whose observed scale heritability (*h*^*2*^) estimates demonstrated significant (*q*<0.05) enrichment of loci in genic and loss-of-function (LoF) intolerant regions of the genome.

| *Trait* | *Genic $h^2$ Fold Enrichment ($p$)* | *LoF Intolerance $h^2$ Fold Enrichment ($p$)* |
| --- | --- | --- |
| Schizophrenia | 1.16 ($5.46 \times 10^{-11}$) | 1.92 ($8.80 \times 10^{-21}$) |
| Educational Attainment | 1.12 ($1.70 \times 10^{-10}$) | 1.79 ($8.65 \times 10^{-23}$) |
| Cognitive Performance | 1.12 ($2.71 \times 10^{-9}$) | 1.79 ($6.26 \times 10^{-29}$) |

### Partitioned Heritability

Heritability partitioning was performed with LDSC using 53 baseline genomic annotations from Finucane, *et al.*^24^ characterizing important molecular properties such as allele frequency distributions, conserved regions of the genome, and regulatory elements. We created additional genome-wide annotations for genic, loss-of-function (LoF) intolerant, positively selected, negatively selected, and Neanderthal-introgressed positions. These evolutionary annotations are described below using per-SNP measurements obtained directly from the original publications. In line with previous studies,^14; 25; 26^ annotations of evolutionary pressures were analyzed as bins of the top 2%, top 1%, and top 0.5% of scores genome-wide. Enrichments were interpreted as follows: (1) all fold-enrichments below zero were considered invalid and not used for any analyses, (2) fold-enrichment values between zero and one indicated *h*^*2*^ depletion attributed to an annotation, and (3) fold-enrichments greater than one indicated *h*^*2*^ enrichment attributed to an annotation.

The genic annotation of the genome was defined by mapping SNPs to genic regions according to the Exome Aggregation Consortium (ExAC) database.^27^ To annotate LoF intolerant regions of the genome, we assigned a probability of loss-of-function (pLI) score to each gene based on the ExAC database. We annotated any SNP in a gene with pLI ≥ 0.9 as LoF intolerant.^14; 27^ Both annotations were coded in a binary manner: genic *versus* not-genic and LoF intolerant *versus* LoF tolerant.

One measure of BGS was tested. The *B-*statistic describes reduction in local allele diversity as a consequence of purifying selection. This measurement leverages phylogenetic information from other closely related species (*e.g.*, gorilla, chimpanzee, orangutan, and rhesus macaque) to test for evidence of pressures selectively removing alleles at a locus from the human genome.^25; 28^ Note that *B* values were transformed (1-*B*) such that *B* near one indicated strong effects of BGS.

We tested *h*^*2*^ enrichment based on three measures of positive selection. Firstly, we examined two estimates of extended haplotype homozygosity (EHH), which quantifies the number of unique haplotypes that exist in a population given a genetic distance. A high EHH score suggests few haplotypes exist in a population, which occurs when an allele is positively selected, dragging along its linked genetic regions. The integrated haplotype score (iHS) incorporates LD and the ancestral *versus* derived context of an allele into EHH. Extreme iHS scores signify a difference between individuals with the ancestral and derived versions of the allele. This test probes for ancient selection trends in the context of selection on a particular genotype at a locus. The cross-population extended haplotype homozygosity (XP-EHH) measure compares haplotypes across populations (here we used 1000 Genomes Project African and European super populations),^29^ rather than across ancestral and derived groupings, to detect instances where one allele has reached fixation in one population but remains polymorphic in the other population. XP-EHH provides evidence of more recent positive selection in the time since the two selected populations diverged.^29^ Finally, the composite of multiple signals (CMS) uses a combination of three measures of selection (long haplotypes, differentiated alleles, and high frequency derived alleles) to detect selective events occurring approximately 20-35 thousand years ago.^30; 31^

Like the haplotype-based estimates of positive selection, Neanderthal local ancestry (LA) is quantified by comparing Neanderthal and contemporary human genomes for human genomic positions with high likelihood of originating via admixture events with Neanderthals since the time that these populations diverged. Enrichment of loci with Neanderthal LA in human complex traits would indicate significant contribution of Neanderthal ancestry to those phenotypes.^32; 33^

Annotations also were generated for summary data describing genotype networks (*i.e.*, a collection of genotypes and their relatedness) across the genome.^34; 35^ The attributes of genotype networks used here are: the number of vertices, average path length, number of components, and the average degree of genotypes or network nodes. The number of vertices and average path length of a genotype network describe how a network extends through genotype space and how heterogeneous or genetically distant the genotypes in a population are. Specifically, the number of vertices is equivalent to the number of distinct genotypes in a population. The average path length describes the average of all possible shortest paths between pairs of genotypes within a network. Average path lengths therefore correspond to the number of SNP changes required to move from one node to another. The remaining properties of genotype networks (number of components and average degree of genotypes) have been used to describe the robustness of a locus to mutational events. The number of components in a genotype network is the number of subgroups in a network connected through a continuous path of edges connecting network nodes. Lastly, the average degree of a network is the average degree of all nodes within that network such that nodes without edges are considered “isolates” and have degree zero. All nodes, including degree zero nodes, are included in the calculation of average degree per genotype network. Genotype networks can be studied to identify the sensitivity and survivability of a phenotype given cycles of neutral mutational events.^36^ That is, populations with similar phenotypes accumulated neutral or slightly deleterious mutations followed by a beneficial mutation that increases population fitness. This type of event fosters development of new genotypes which lie along a similar genotype network (described by the features detailed above). Enrichment of genotype networks may demonstrate the ability of a phenotype to tolerate a population’s exploration of genotype space over time and the chance that it tolerates neutral and/or deleterious events in favor of future beneficial evolutionary events.^36^

### Effect Size Distribution

GWAS have detected tens-to-hundreds of loci associated with complex traits but individual locus effect sizes are relatively small. Studying these loci in combination explains substantially greater trait heritability than any individual locus. Descriptive statistics of the effect size distribution for loci associated with complex traits were the main dependent variable of interest in this study. We evaluate how evolutionary pressures (described above) contribute to the descriptive statistics of trait associated loci effect size distribution described by Zhang*, et al.*^2^

The GENESIS (GENetic Effect-Size distribution Inferences from Summary-level data) R package was used to determine effect size distribution descriptive statistics for brain-related traits.^2^ All GWAS for brain-related traits were filtered to include only Hapmap3 SNPs.^37^ As per developer guidelines,^2^ SNPs were removed if (1) their effective sample sizes were less than 0.67 times the 90^th^ percentile of the per-SNP sample size distribution, (2) they fell within the major histocompatibility region (these were removed due to the complex LD structure of this region), and (3) they had extremely large effect sizes (per-SNP effect *z*-scores>80). Descriptive statistics of effect size distribution are defined as follows: (1) π_c_ is the proportion of susceptibility SNPs per trait, (2) σ^2^ is the variance parameter for non-null SNPs, and (3) *a* is the parameter describing all residual effects not captured by the variance of effect-sizes (*e.g.*, population stratification, underestimated effects of extremely small effect size SNPs, and/or genomic deflation).^2^ Zhang, *et al*.^2^ compared two- and three-component models of effect size distribution assuming 99% of SNPs in complex trait GWAS are null and the effect sizes for the remaining 1% of non-null SNPs follow a normal distribution centered around zero (two-component) or a mixture normal distribution (three-component). Results from Zhang, *et al.*^2^ support the use of two-component models for *autism spectrum disorder*, *bipolar disorder*, *major depressive disorder*, *schizophrenia*, *college completion*, *neuroticism*, *cognitive performance*, and *intelligence quotient*. We therefore considered only the two-component model to calculate effect size distribution descriptive statistics for the brain-related phenotypes analyzed in this study.

### Heterogeneity Features

Many psychiatric disorder diagnoses rely on the number of diagnostic criteria per patient exceeding some threshold. Heterogeneity arises when individuals with the same diagnosis exhibit different combinations of diagnostic criteria. Here we used the psychiatric disorder heterogeneity features described by Olbert, *et al.*^20^ to evaluate the impact of psychiatric disorder diagnostic heterogeneity on effect size distribution descriptive statistics for GWAS of psychiatric disorders. We focus our analyses on psychiatric disorder *total symptoms* and *diagnostic combinations*. *Total symptoms* are the number of criteria considered for diagnosis according to DSM-5. *Diagnostic combinations* were defined as any set of symptoms that could meet diagnostic criteria for a disorder dependent upon the presence of a certain combination of symptoms. Based on DSM-5 descriptions of these measures, *obsessive compulsive disorder* and *Tourette syndrome*^38^ would be characterized by the same *total symptoms* and *diagnostic combinations*, so only *Tourette syndrome* was included in regression analyses because it has a higher *h*^*2*^ *z*-score than *obsessive compulsive disorder*.

### Non-Parametric Correlation and Regression Tests

Spearman’s rho (ρ) is a non-parametric test that measures the strength of association between two variables. We used ρ to test the correlation between: (i) standardized (*e.g., z*-score-converted) dependent (*e.g.*, trait *h*^*2*^ and effect size distribution descriptive statistics) and independent measures (*e.g.*, enrichments of genic and LoF-intolerant loci and loci demonstrating evidence of natural selective pressures and Neanderthal introgression), (ii) standardized measures of evolutionary pressure and observed-scale *h*^*2*^ and unstandardized measures of effect size distribution, and (iii) unstandardized measures of all evolutionary pressures, observed-scale *h*^*2*^, and measures of effect size distribution. Unless otherwise noted, we discuss results using standardized independent and dependent variables but provide all correlations in the corresponding supplementary information. Non-parametric regression tests included median-based linear regression (MBLM; y ~ x), local regression (locally-estimated scatterplot smoothing or LOESS: y ~ x_1_ + x_2_ + x_n_), and generalized additive models (GAM: y ~ x_1_ + x_2_ + x_n_ and y ~ x_1_ × x_2_ × x_n_) to evaluate the predictive capabilities of linear and additive models on an outcome variable of interest. MBLM computes all lines between each pair of data points and reports the median of the slopes of these lines. Model *r*^*2*^ values are not reported due to the median-based nature of the test; however, MBLM slopes and *p*-values for each slope are reported. GAMs were evaluated using nominally significant MBLM predictors. When multiple binary versions of the same genomic annotation were significant predictors of the outcome variable by MBLM (*e.g.*, enrichment of loci in the top 2%, 1%, and 0.5% of *B-*statistics were all significant predictors of outcome Y), the genomic annotation with the lowest *p*-value was included in GAM. GAMs are accompanied by generalized cross-validation (GCV) scores representing mean square error of the model. GCVs associated with GAM estimate the model prediction error without performing formal cross-validation in an out-sample. GCVs closer to zero represent better fitting models. The analysis of variance (ANOVA; `anova.gam`) feature in the `mgcv`^39^ R package was used to evaluate significant differences in model fit.

### Multiple Testing Correction

We applied a false discovery rate (FDR) multiple testing correction for the expected proportion of false discoveries among rejected null hypotheses based on the number of tests performed in each analysis. Conversion of test statistic *p*-values to multiple testing corrected *q*-values was performed in R using the `p.adjust` function and the `fdr` method.^40^ Unless otherwise noted, we report original, unadjusted p-values for enrichment estimates, correlations, and model estimates. For example, `anova.gam` prints pre-adjusted test statistics; we report these as *q* instead of *p*.

## Results

A total of 75 mental health outcomes were sufficiently heritable (*h2 z*-score ≥ 7) for partitioning based on evolutionarily relevant functional annotations of the genome (Table S1). These phenotypes were grouped into eleven categories based on general trait similarities: *anxiety and related*, *brain imaging*, *cognition*, *cross disorder*, *depressive and related*, *eating*, *externalizing behavior*, *family adversity*, *internalizing behavior, neurodevelopmental*, and *substance use and related* (Table S1).

### Enrichment of Genic and Loss-of-Function Intolerance

The *h*^*2*^ estimates of 39 and 69 phenotypes out of the 75 evaluated were nominally significantly enriched for genic and LoF-intolerant loci, respectively, 34 and 67 of which survived FDR multiple testing correction (*q*<0.05; Figure 1 and Table S1). The top three significant phenotypes were the same for both annotations: *schizophrenia*, *educational attainment*, and *cognitive performance* (Table 1). The greatest *h*^*2*^ enrichment of genic loci was observed in the GWAS of *T1-weighted left-plus-right caudate volume* (enrichment=1.32-fold, *p*=0.003). When phenotypes were binned into eleven major categorical domains (Table S1), the *brain imaging* category demonstrated significantly higher enrichment of *h*^*2*^ attributed to genic loci than *cognition* (difference in mean=0.174, *q*=1.47×10^−5^), *depressive and related* (difference in mean=0.125, *q*=0.002), *internalizing behavior* (difference in mean=0.130, *q*=0.001), and *substance use and related* (difference in mean=0.158, *q*=1.46×10^−5^) phenotype categories. The *h*^*2*^ for GWAS of *confiding adult relationship* (enrichment=2.31-fold, *p*=1.16×10^−4^) and *recent easy annoyance* (enrichment=2.31-fold, *p*=1.60×10^−5^) demonstrated the greatest LoF-intolerance enrichment magnitude. There was no significant difference in enrichment of LoF intolerance loci by phenotype category (ANOVA *p*=0.149). Thirty-two phenotypes demonstrated FDR significant enrichment of both genic and LoF intolerant loci (*q*<0.05); there was a strong overlap between genic and LoF intolerant enrichment magnitudes (ρ=0.490, *p*=0.003; Figure S1).

**Figure 1.**
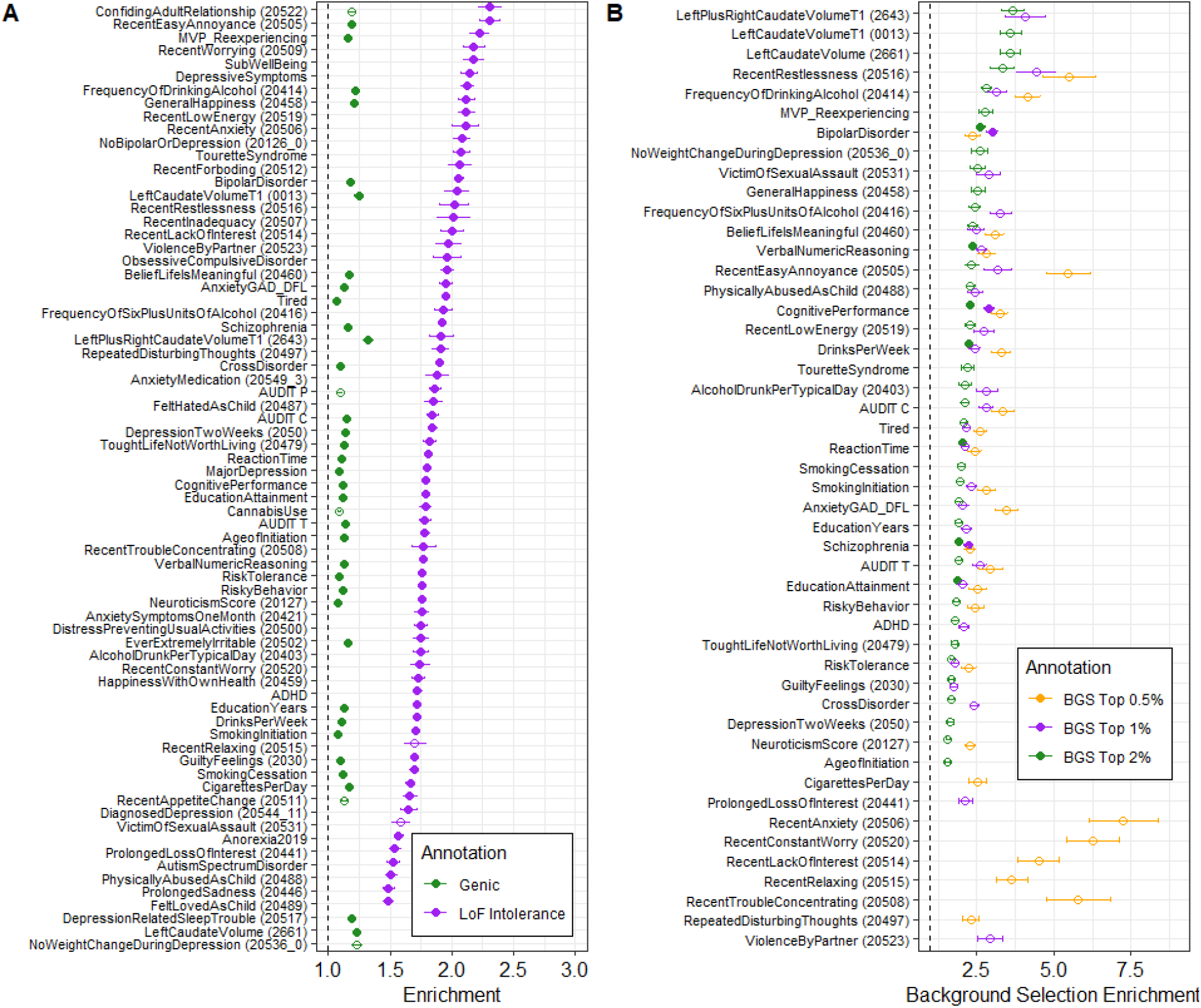
Enrichment of natural selection and functional annotation measures. Significant enrichments of genic and loss-of-function (LoF) intolerant loci (A) and three genomic annotations of background selection (BGS). Both panels list phenotypes in descending order from highest to lowest magnitude enrichment for the most abundant annotation (*i.e.*, LoF intolerance in panel A and top 2% of BGS scores in panel B). All phenotypes listed are at least nominally significant (*p*<0.05) and solid circles indicate that the enrichment survived multiple testing correction (*q*<0.05). Error bars represent the 95% confidence interval around each enrichment estimate.

We next tested whether significant genic and LoF intolerance enrichments were influenced by trait heritability as a proxy representation of effective population size per phenotype. Standardized genic (ρ=0.950, *p*=2.20×10^−16^) and LoF-intolerant (ρ=0.890, *p*=2.20×10^−16^) enrichments were significantly positively correlated with standardized trait *h*^*2*^ (Figure S1). These results are recapitulated in unstandardized correlations where enrichment magnitude for genic and LoF-intolerance annotations was significantly negatively correlated with trait standardized *h^2^ z*-score (standardized *h*^*2*^ versus genic enrichment: ρ=-0.790, *p*=3.52×10^−10^; standardized *h*^*2*^ versus LoF intolerance enrichment: ρ=-0.300, *p*=0.015; Figure S1). In other words, GWAS for phenotypes with higher confidence *h*^*2*^ estimates tend to show higher confidence enrichment estimates, but these estimates tend to approach 1 (*i.e.*, minimal enrichment). The former observation is expected and suggests that the more accurately trait *h*^*2*^ is estimated, the more accurately that *h*^*2*^ can be partitioned.^24^ The latter observation is perhaps unexpected and may suggest that the magnitude of *h*^*2*^ enrichment attributed to functional annotation is nonuniformly a function of proxy measures of sample size up to a certain level of trait *h*^*2*^, visually estimated here to be an approximate *h^2^ z*-score = 15.

### Enrichment of Background Selection

The GWAS of 48 phenotypes demonstrated nominally significant enrichment of BGS in at least one of the genomic annotations created here (*i.e.*, top 2%, 1%, and/or 0.5% of *B-*statistic genome-wide). The only major phenotype domain not demonstrating enrichment of BGS was *eating*, comprising the *anorexia nervosa* and *recent appetite change* phenotypes. Elevated BGS was detected at the level of FDR significance (*q*<0.05) in the GWAS of seven phenotypes, all of which demonstrated at least nominally significant BGS enrichment at all three genomic annotations and when using the *B-*statistic as a continuous annotation (Table 2). Three of these phenotypes also demonstrated depletion of positive selective pressures at various levels of thresholding each annotation of the genome: (1) the *h*^*2*^ of *cognitive performance* was depleted of SNPs in the top 2% of CMS scores (0.112-fold depletion, *p*=0.002) and SNPs in the top 1% of iHS scores (0.207-fold depletion, *p*=0.012), (2) *schizophrenia* was depleted of SNPs in the top 2% of XP-EHH scores (0.216-fold depletion, *p*=0.005), and (3) *verbal numeric reasoning* was depleted for SNPs in the top 2% of XP-EHH scores (0.190-fold depletion, *p*=0.023), SNPs in the top 2% of CMS scores (0.101-fold depletion, *p*=0.028), and SNPs in the top 2% of iHS scores (0.371-fold depletion, *p*=0.001). Table S1 lists enrichment estimates and *p*-values for all three annotations of each selection statistic (BGS, iHS, XP-EHH, and CMS). Because BGS results in reduced genetic variation, these enrichment results for BGS and positive selection may be interpreted as concordant evidence for the strong effect of BGS on complex traits.

**Table 2.** Background selection enrichments. Seven phenotypes whose observed scale heritability (*h*^*2*^) estimates demonstrated significant enrichment (*q*<0.05) of background selection (BGS) in at least one genomic annotation of per-SNP *B-*statistic measures. FDR significant observations (*q*<0.05) are in bold text.

| *Trait* | *BGS (top 2%) $h^2$ Fold Enrichment ($p$)* | *BGS (top 1%) $h^2$ Fold Enrichment ($p$)* | *BGS (top 0.5%) $h^2$ Fold Enrichment ($p$)* |
| --- | --- | --- | --- |
| Bipolar Disorder | **2.62 ($2.01 \times 10^{-6}$)** | **3.01 ($4.31 \times 10^{-6}$)** | 2.36 (0.021) |
| Cognitive Performance | **2.29 ($1.81 \times 10^{-8}$)** | **2.90 ($7.31 \times 10^{-6}$)** | 3.24 (0.001) |
| Drinks Per Week | **2.25 ($8.33 \times 10^{-7}$)** | 2.43 (0.004) | 3.29 (0.006) |
| Educational Attainment | **1.86 ($9.69 \times 10^{-6}$)** | 2.04 (0.001) | 2.53 (0.003) |
| Schizophrenia | **1.90 ($7.29 \times 10^{-6}$)** | **2.23 ($6.04 \times 10^{-6}$)** | 2.27 (0.007) |
| Reaction Time | **2.02 ($1.33 \times 10^{-7}$)** | 2.13 (0.001) | 2.44 (0.018) |
| Verbal Numeric Reasoning | **2.35 ($8.63 \times 10^{-8}$)** | 2.63 ($2.21 \times 10^{-4}$) | 2.81 (0.017) |

We next tested whether standardized BGS enrichments were associated with standardized *h*^*2*^ (like those observations with genic and LoF intolerant loci). There were significant positive associations between standardized *h*^*2*^ and BGS at all three genomic partitions: *h*^*2*^ versus BGS_Top2%_: ρ=0.911, *p*=2.20×10^−16^, *h*_*2*_ versus BGS_Top1%_: ρ=0.856, *p*=1.7×10^−9^, *h*_*2*_ versus BGS_Top0.5%_: ρ=0.751, *p*=9.81×10^−6^. Unstandardized correlations reiterate observations with genic and LoF intolerance correlations where those phenotypes with high confidence *h*^*2*^ estimates (*e.g., h^2^ z*-score greater than ~15) generally showed no relationship between *h*^*2*^ and BGS enrichment (Figure S2).

### Enrichment of Neanderthal Introgression

Posterior probability of Neanderthal LA indicates contribution of Neanderthal genomes to a particular phenotype.^33; 41; 42^ The observed scale *h*^*2*^ of one phenotype (UK Biobank *neuroticism score*) was significantly depleted of SNPs in the top 2% of Neanderthal LA 0.358-fold depletion, *p*=8.61×10^−6^). This phenotype also exhibited nominally significant enrichment of SNPs demonstrating evidence of BGS in the top 2% of *B*-statistic (1.57-fold enrichment, *p*=0.003) and top 0.5% of *B*-statistic (2.28-fold enrichment, *p*=0.002). It has been shown that per-SNP measures of Neanderthal LA and *B*-statistic are significantly correlated^33^ however we demonstrate, with the *τc* statistic calculated in LDSC,^14; 24^ that enrichment of Neanderthal LA in the UKB phenotype *neuroticism score* was independent of all other annotations of the genome investigated in this study (*p*=1.41×10^−4^).^41^

### Correlates of Effect Size Distribution

Parameters describing effect size distribution per phenotype (π_c_, σ^2^, and *a*) were generated using the GENESIS R package (Figure 2).^2^ Three effect size distribution outliers were detected with measurements of at least three standard deviations from the mean. First, *left plus right caudate volume* (*z*-score=3.06) was an outlier with respect to the σ^2^ parameter, highlighting a wide distribution of variant effect sizes underlying this phenotype. Second, *anorexia nervosa* (*z*-score=3.75) and *bipolar disorder* (*z*-score=5.78) were outliers with respect to the *a* parameter, suggesting possible residual population stratification underlying these GWAS. Notably, neither phenotype is an outlier with respect to their intercept from LDSC: *anorexia nervosa intercept*=1.03±0.010 and *bipolar disorder* intercept=1.01±0.008.

**Figure 2.**
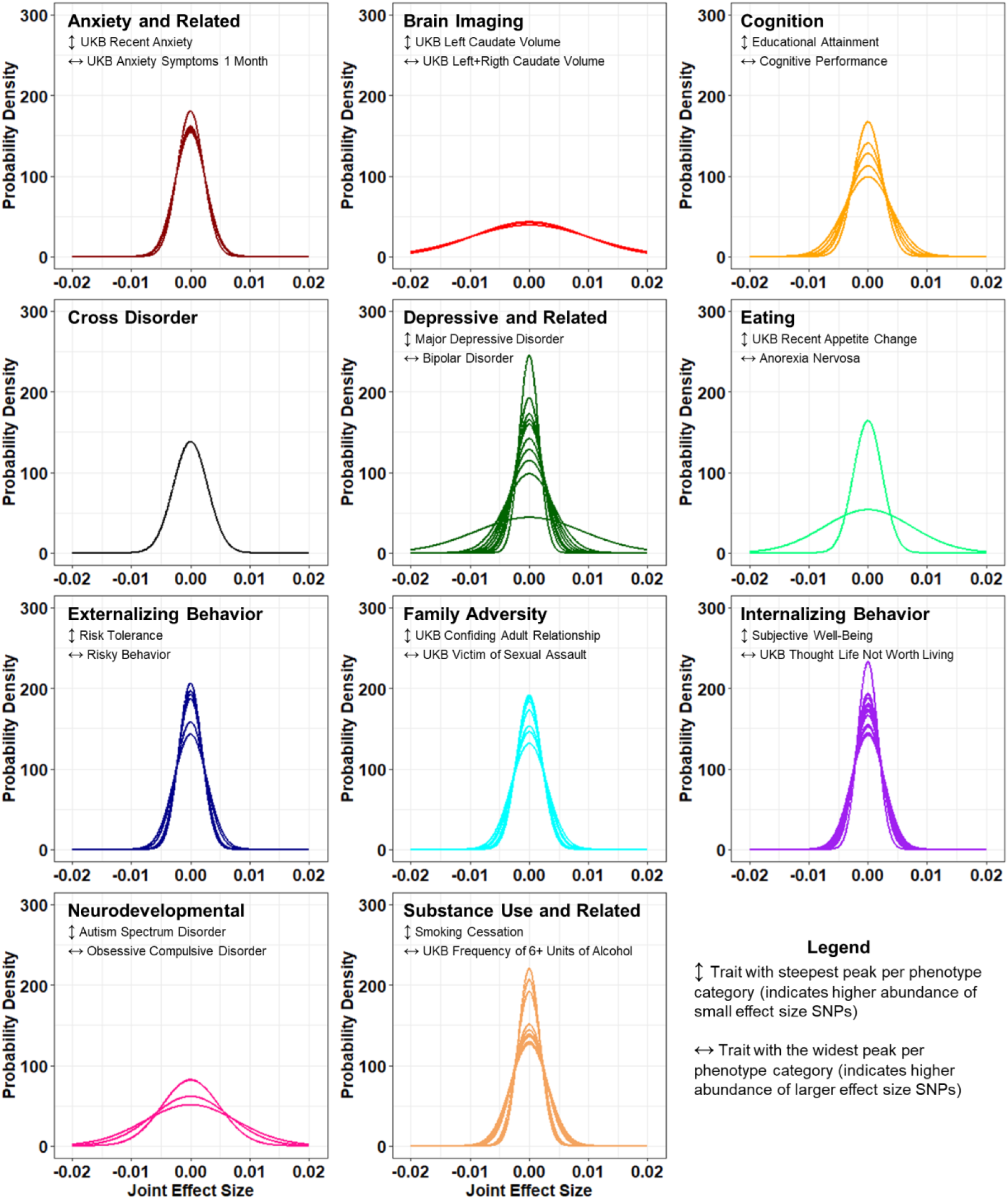
Complex trait effect size distribution curves. Effect size distribution curves derived from GWAS summary statistics for 75 mental health and disease phenotypes using the GENESIS R package. Phenotypes are grouped into eleven categories (Table S1). The phenotype corresponding to the steepest curve (↕) and the widest curve (↔) are identified in each subplot.

We used the non-parametric Spearman correlation test to evaluate linear relationships between effect size distribution parameters (π_c_, σ^2^, and *a*) and (1) BGS, (2) genic and LoF intolerance, (3) trait *h*^*2*^ (considering liability-scale *h*^*2*^ for binary traits), and (4) diagnostic heterogeneity (psychiatric disorders only). The phenotype attributes *total symptoms* and *diagnostic combinations* were obtained from Olbert, *et al.* 2014 and describe how the total number of DSM-5 criteria per psychiatric disorder contributes to diagnostic heterogeneity.^20^ There were no statistically significant differences in correlation when outliers were removed so all reported correlations included the complete sample. Standardized genic and LoF-intolerance enrichments significantly predicted all three effect size parameters (ρ_πc_genic_=0.851, *p*=2.20×10^−16^; ρ_πc_LoF_=0.600, *p*=4.68×10^−7^; ρ_σ2_genic_=0.862, *p*=2.20×10^−16^; ρ_σ2_LoF_=0.599, *p*=5.07×10^−7^; ρ_*a*_genic_=0.798, *p*=5.72×10^−9^; ρ_*a*_LoF_=0.736, *p*=2.20×10^−16^; Figure 3). Using unstandardized measures of genic and LoF intolerance enrichment, we observed no relationship between LoF intolerance and effect size distribution; however, enrichment of genic loci was negatively correlated with π_c_ (ρ_πc_genic_=-0.682, *p*=1.52×10^−6^) and positively correlated with σ^2^ (ρ_σ2 genic_=0.546, *p*=2.33×10^−4^). This suggests that *h*_*2*_ enrichment attributed to loci in genic regions of the genome (1) decreases the number of detectable risk loci and (2) increases the variance associated with effect size estimates for those risk loci that can be detected.

**Figure 3.**
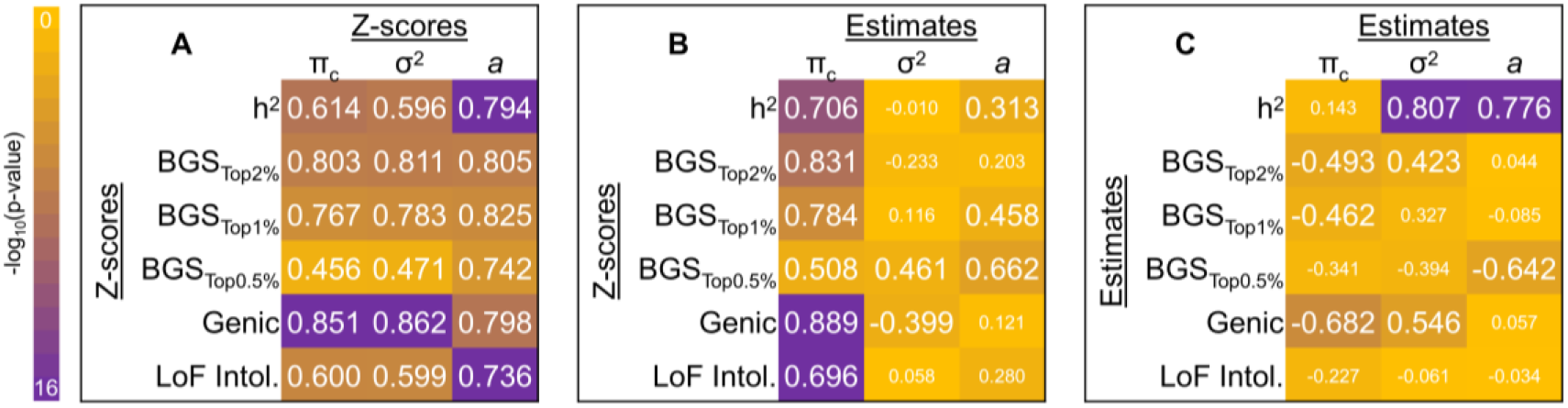
Correlation between effect size distribution, natural selection, and functional annotation. Spearman correlation between effect size distribution descriptive statistics and heritability (*h*^*2*^), enrichment for background selection (BGS) partitions (top 2%, 1%, and 0.5%), genic loci, and loss-of-function intolerant (LoF Intol.) loci. Panel A correlates *z*-scores for all measures, panel B correlates the *z*-score for all measures against unstandardized effect size distribution descriptive statistics, panel C correlates unstandardized enrichment and *h*^*2*^ magnitudes against unstandardized effect size distribution descriptive statistics. Large text indicates significant correlation after multiple testing correction (*q*<0.05).

Using only phenotypes with at least nominal enrichment of BGS, all three standardized effect size distribution parameters were nominally correlated with standardized BGS: π_c_ (ρ_πc_Btop2%_=0.557, p=3.59×10^−7^; ρ_πc_Btop1%_=0.557, *p*=5.49×10^−7^; ρ_πc_Btop0.5%_=0.575, *p*=1.26×10^−7^), σ^2^ (ρ_σ2_ Btop2%_=0.514, *p*=3.39×10^−6^; ρ_σ2_ Btop1%_=0.549, *p*=5.16×10^−7^; ρ_σ2_ Btop0.5%_=0.613, *p*=9.80×10^−9^), and *a* (ρ_*a*_ Btop2%_=0.688, *p*=2.20×10^−16^; ρ_*a*_ Btop1%_=0.632, *p*=9.18×10^−10^; ρ_*a*_ Btop0.5%_=0.543, *p*=7.49×10^−7^; Figure 3). Unstandardized analyses reveal that higher enrichment of BGS generally associated with reduced proportion of risk loci (π_c_; ρ_πc_Btop2%_=-0.493, *p*=0.002; ρ_πc_Btop1%_=-0.462, *p*=0.012), increased variance among those non-null risk loci (σ^2^; ρ_σ2_ Btop2%_=0.413, *p*=0.009), and decreased residual effects (*e.g.*, population stratification, underestimated effects of extremely small effect size SNPs, and/or genomic deflation) as measured by the *a* parameter (ρ_*a*_ Btop0.5%_=-0.642, *p*=5.50×10^−4^). These results highlight the relative importance of BGS on detection of genetic risk for complex phenotypes. Considering psychiatric diagnoses only, there were no detectable correlative relationships between measures of heterogeneity (total symptoms and diagnostic combinations) and effect size distribution (π_c_, σ^2^, and *a*).

### Predicting Effect Size Distributions

We next tested the ability of each unstandardized trait property (BGS, genic and LoF intolerance enrichment, and heterogeneity features) to predict effect size distribution descriptive statistics (π_c_, σ^2^, and *a*) using median-based linear models (MBLM) (Figure 4). For this analysis, Spearman correlations were not used to inform MBLM testing because of potential effects of (1) confounder bias, (2) collider bias, and/or (3) incidental cancellation potentially resulting in (a) a correlated independent variable not predicting an effect size distribution descriptive statistic or (b) an uncorrelated independent variable predicting an effect size distribution descriptive statistic. Therefore, enrichment of genic, LoF intolerant, and all partitions of BGS were used in MBLM to predict π_c_, σ^2^, and *a* regardless of their pairwise correlations. Though highly correlated with effect size distributions (Figures S1 and S2), standardized and unstandardized trait *h*^*2*^ was not used to predict effect size distribution parameters because of its representation of effective population size in case-control phenotypes which varies considerably across phenotypes.

**Figure 4.**
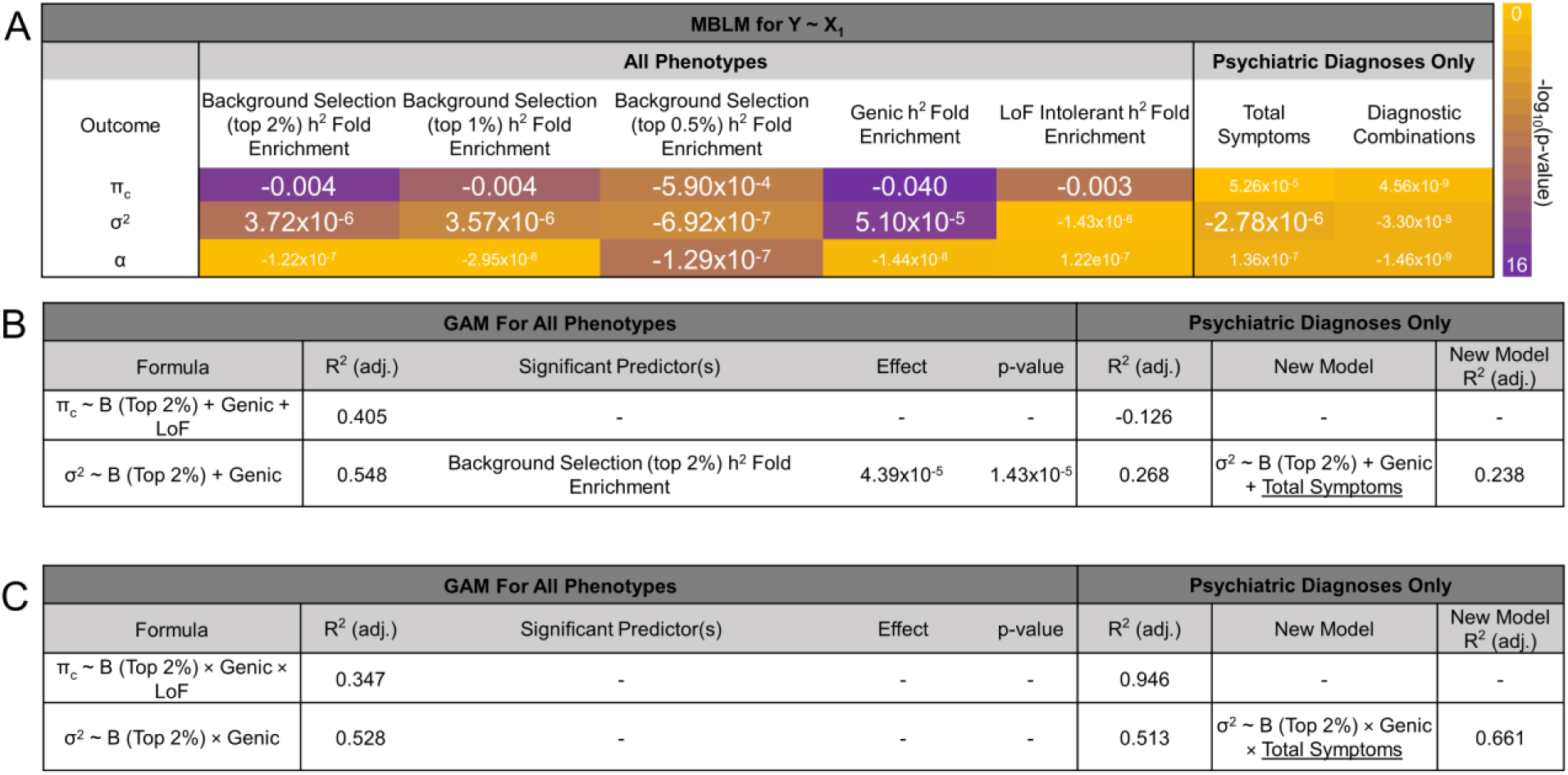
Predicting effect size distribution with natural selection, functional annotation, and phenotype heterogeneity. Summary of models predicting effect size distribution parameters. (A) Median-based linear models (MBLM) for predicting effect size distribution descriptive statistics with a single unstandardized independent variable. Larger font indicates at least nominal significance for the respective effect of each independent variable on the outcome effect size distribution descriptive statistic. (B and C) Generalized (GAM) additive (B) and interactive (C) models of effect size distribution descriptive statistics using individually significant predictors of each metric (from A). Prior to testing the addition of heterogeneity information, each model was re-evaluated in psychiatric disorders only.

After multiple testing correction, the π_c_ parameter (*i.e.*, proportion of susceptibility loci) was predicted by enrichment of genic, LoF intolerant, and all three BGS partitions (Figure 4). After multiple testing correction, the σ^2^ parameter (*i.e.*, the variance parameter for non-null SNPs) was predicted independently by enrichment of all three BGS partitions and genic loci. In the additive model of σ^2^, enrichment of *h*^*2*^ attributed to loci in the top 2% of genome-wide *B-*statistic was the only significant predictor (β=4.39×10^−4^, *p*=1.43×10^−5^; full model *r*^*2*^=0.548). In the absence of out-sample cross-validation, GCV estimates associated with model fit (*r*^*2*^) represent a prediction error estimate whereby smaller GCV values indicate greater model fit. For the additive model of σ^2^, we observed a very small prediction error in the model fit, GCV=5.79×10^−10^. Interactive models of π_c_ and σ^2^ generally explained comparable variances in the outcome variable of interest but lack significant individual or interactive predictors: π_c_ ~ B_Top2%_ × genic × LoF full model *r*_*2*_=0.347, GCV=2.72×10^−5^ and σ^2^ ~ B_Top2%_ × genic full model *r*^*2*^=0.528, GCV=4.80×10^−10^. The *a* parameter (*i.e.*, residual effects not captured by the variance of effect-sizes) was only predicted by *h*^*2*^ enrichment attributed to loci in the top 0.5% of *B-*statistic scores (MBLM β=-1.29×10^−7^, *p*=2.48×10^−5^) and therefore additive and interactive models were not tested.

We next focused on a subset of phenotypes for which diagnostic heterogeneity information exists (*ADHD*,^43^ *anorexia nervosa*,^44^ *autism spectrum disorder*,^45^ *bipolar disorder*,^46^ *major depressive disorder*,^47^ *schizophrenia*,^48^ *anxiety*,^49^ and *Tourette syndrome*;^38^ Figure 4 and Table S1). With these phenotypes we tested whether diagnostic information such as *number of diagnostic combinations* and *total symptoms* contribute to risk locus effect size distribution descriptive statistics. First, we re-evaluated the original additive models (see above) in this subset and recapitulated the relationship between the σ^2^ parameter, BGS, and genic loci (model *r*^*2*^=0.268, GCV=7.58×10^−10^) but did not identify any independent significant predictors of σ^2^. The additive model for the effects of BGS, genic, and LoF intolerance enrichments on π_c_ were discordant when applied to psychiatric disorders only (model *r*^*2*^=-0.126, GCV=1.89×10^−5^) and again no significant individual predictors were identified. Conversely, the interactive models exhibited comparable (or increased) and concordant model fit in the subset of psychiatric disorders (π_c_ *r*_*2*_=0.946, GCV=1.01×10^−4^ and σ^2^ *r*_*2*_=0.513, GCV=4.80×10^−10^). Next, we included measures of phenotype heterogeneity into the additive and interactive models. The only heterogeneity parameter suggesting putative prediction potential was the number of total symptoms per psychiatric disorder (MBLM β_σ2_total_symptoms_=-2.78×10^−6^, *p*=0.043). When added to the σ^2^ additive model, model fit modestly decreased (*r*^*2*^=0.238, GCV=7.69×10^−10^, *p*_difference_=0.411) with no significant contributors, suggesting no greater prediction potential than the original additive model. Including the total symptoms measure into the interactive model of σ^2^ modestly increased model fit: σ^2^ ~ B (Top 2%) × Genic × Total Symptoms; model *r*^*2*^=0.661, GCV=1.82×10^−9^ (*p*_difference_=0.401).

## Discussion

The effect size distribution of a phenotype may be used to infer the total sample size required for risk locus detection as well as the associated genetic variance explained.^2^ Mental health and disease phenotypes typically demonstrate narrow effect size distributions (*i.e.*, normal distributions with most data points distributed tightly around zero) supporting that they are (1) highly polygenic and (2) the additive result of SNPs conferring small effects.^2^ Here we investigated how these effect sizes and their distribution are shaped by natural selection.

Genetic risk for *schizophrenia*, *blood pressure*, *body mass index*, *major depressive disorder*, and others is influenced by natural selection.^13; 14; 16; 50^ The concept of genome-wide flattening was used to demonstrate that low-frequency variants contribute less to the polygenicity of a trait than common variants, and that negative selection constrains effect sizes for these common variants.^13^ Flattening describes the reduced effect size distributions for complex traits subjected to BGS, resulting in a highly polygenic architecture consisting of many loci with relatively small effects. Using common genetic variation and three separate measures of effect size distributions describing (1) the number of non-null risk loci, (2) the variance of non-null risk loci effect size estimates, and (3) parameter representing residual variance not attributed to non-null risk loci, we converge on comparable findings and replicate the detection of BGS influencing *schizophrenia* risk.^14^ We additionally replicate that the genetic risk loci for *schizophrenia* confer extremely small effects on the phenotype due to the selective removal of larger effect variants from the genome over evolutionary time. Here, we extend this observation to 48 other mental health and disease phenotypes and identify traits whose risk loci exhibit an overrepresentation of BGS. Seven phenotypes (*bipolar disorder*, *cognitive performance*, *drinks per week*, *educational attainment*, *schizophrenia*, *reaction time*, and *verbal numeric reasoning*) survived multiple testing correction, suggesting a significant influence of BGS on their common genetic variation. We also detect an overabundance of genic and LoF intolerant loci contributing to phenotype *h*^*2*^, implying that functionally important regions of the genome contribute the most to *h*^*2*^.^13; 14; 27; 51^

Three phenotypes demonstrated enrichment of BGS, genic, and LoF intolerance: *cognitive performance*, *educational attainment*, and *schizophrenia*. These overlapping phenotypes demonstrate negatively genetically correlated and arguably advantageous (*cognitive performance* and *educational attainment*) and disadvantageous (*schizophrenia*) features of the mental health phenome, potentially due to evolutionary tradeoffs associated with similar biological effects. The persistence of risk-conferring (*i.e.*, deleterious) loci in regions of the genome intolerant to mutation and at common frequency may at first be viewed as paradoxical. We and others hypothesize that this paradox could be attributed to BGS, which selectively removes haplotypes from the population that contain large effect deleterious mutations.^52; 53^ This in turn reduces genetic diversity and allows for small-effect variants (or haplotypes) to rise in frequency to the levels of common variation detected by GWAS of the mental health and disease phenome.^14; 27; 52–54^

We next evaluated the ability of BGS, genic, and LoF intolerance enrichments to predict the effect size distributions of the mental health and disease phenome. We identified two models for predicting the variance parameter (σ^2^) of effect size distributions using BGS and genic locus enrichment: (1) an additive model that explained 54.8% of the variance in σ^2^ and (2) an interactive model that explained 52.8% of the variance in σ^2^. These data explicitly quantify the effects of natural selection on the distribution of effects sizes underlying GWAS of complex traits. However, they do not suggest that BGS acts directly on the analyzed phenotypes. They may instead point to strong effects of BGS on pleiotropic regions of the genome shared by these phenotypes.^12^ Only the interactive model performed modestly well when tested in psychiatric disorders only, explaining 51.3% of the variance in σ^2^. After incorporating phenotype heterogeneity information, 66.1% of variance in the σ^2^ parameter (*i.e.*, the variance in effects sizes for non-null risk loci) for psychiatric disorders was explained by the interaction between BGS, loci in genic regions of the genome, and the total number of symptoms included in psychiatric disorder diagnosis, though this improvement over the model excluding total symptom count was not significant.

Though increased BGS and genic locus enrichment produce greater σ^2^ estimates (greater effect size variance, implying some relatively large and some relatively small effect risk loci), increased phenotype heterogeneity from total symptom counts may reduce the potential for discovering (1) greater numbers of risk loci and (2) higher risk conferring loci. This is likely especially true for traits like *ADHD* and *PTSD* which have substantially greater symptom counts (which can be considered as different phenotypically and of which only a subset are required for diagnosis) than other psychiatric disorders.^20^ These data also lend support to the trends of (1) utilizing continuous phenotype definitions (*e.g.*, GWAS of PTSD Check List (PCL) criterion count^55^) instead of case-control diagnoses, where appropriate, and (2) studying endophenotypes rather than diagnoses (*i.e.*, phenotypes connecting behavioral symptoms or psychopathology with well-understood biological phenotypes; *e.g.*, smooth pursuit eye movements as an intermediate phenotype for schizophrenia).^56–58^

Our study has three primary limitations. First, a model predicting the effect size distribution variance parameter (σ^2^) of psychiatric disorders is inherently limited in the number of observations contributing to the development of said model. The scarcity of data contributing to model training likely drives this model towards overfitting and a lack of generalizability, even via cross-validation. This is a difficult issue to tackle, however, due to (1) the motivation to increase the training set size, (2) decisions of appropriate test set size, and (3) lack of large numbers of psychiatric disorders to test. We do not attempt to cross validate this model due to a lack of data to do so, but we do demonstrate that the original model of interactive effects between genic and BGS loci persisted in psychiatric disorders. Second, two psychiatric disorders (*anorexia nervosa* and *bipolar disorder*) were statistical outliers with respect to *a*, which captures residual variance in effect size estimates. Though not evident by their LDSC intercept estimates, it is possible that these analyses harbor residual population stratification requiring longer LD-blocks to resolve.^59^ Finally, we use enrichment values to make predictions about effect size distribution outcomes. The precise magnitude of enrichment may change depending on (1) LDSC model structure^12; 24; 51^ and, as we demonstrate, (2) robust standardized *h*^*2*^ scores. Therefore, this model will require further refinement as GWAS for complex phenotypes continue to grow.

This study supports the extensive effect of BGS and overrepresentation of genic and LoF-intolerant loci in the GWAS of complex mental health and disease phenotypes. These features were capable of predicting the variance in genetic risk effect sizes for these neuropsychiatric phenotypes. We demonstrate that psychiatric disorder genetic risk effect sizes may be masked by phenotype heterogeneity and elucidate genetic and evolutionary evidence in favor of the continuous and/or intermediate phenotype approaches to studying the genetics of psychiatric disorders and complex phenotypes. These data begin to unravel questions of “how” and “why” human brain-related GWAS suffer a substantial burden from large sample size requirements by demonstrating how the combined effects of functional regions and natural selection influence SNP effect size distributions. This research is supported in part by the Department of Veterans Affairs Office of Academic Affiliations Advanced Fellowship Program in Mental Illness Research and Treatment, the Department of Veterans Affairs National Center for Post-Traumatic Stress Disorder Clinical Neurosciences Division, and the VA Connecticut Healthcare System. The views expressed here are the authors’ and do not necessarily represent the views of the Department of Veterans Affairs.

## Supplemental Data

Supplemental data include one table and two figures.

## Acknowledgements

This study was supported by the Simons Foundation Autism Research Initiative (SFARI Explorer Award: 534858), the American Foundation for Suicide Prevention (YIG-1-109-16), the National Institutes of Health (R21 DC018098 and R21 DA047527), and the National Center for PTSD of the U.S. Department of Veterans Affairs. This research is supported in part by the Department of Veterans Affairs Office of Academic Affiliations Advanced Fellowship Program in Mental Illness Research and Treatment, the Department of Veterans Affairs National Center for Post-Traumatic Stress Disorder Clinical Neurosciences Division, and the VA Connecticut Healthcare System. The views expressed here are the authors’ and do not necessarily represent the views of the Department of Veterans Affairs.

## Declaration of Interests

The authors declare no competing interests.

## Web Resources

PGC data download: https://www.med.unc.edu/pgc/download-results/; SSGAC data download: https://www.thessgac.org/data; UKB data showcase: http://biobank.ndph.ox.ac.uk/showcase/; UKB Brain Imaging Genetics server – version 2.0: http://big.stats.ox.ac.uk/

## Supplementary Information

**Table S1.** Description of all phenotypes, data sources, and statistics derived from each GWAS of the mental health and disease phenome. Nominally significant enrichments are provided; bolded enrichments survive multiple testing correction (*q*<0.05) [xlsx file attached].

**Figure S1.**
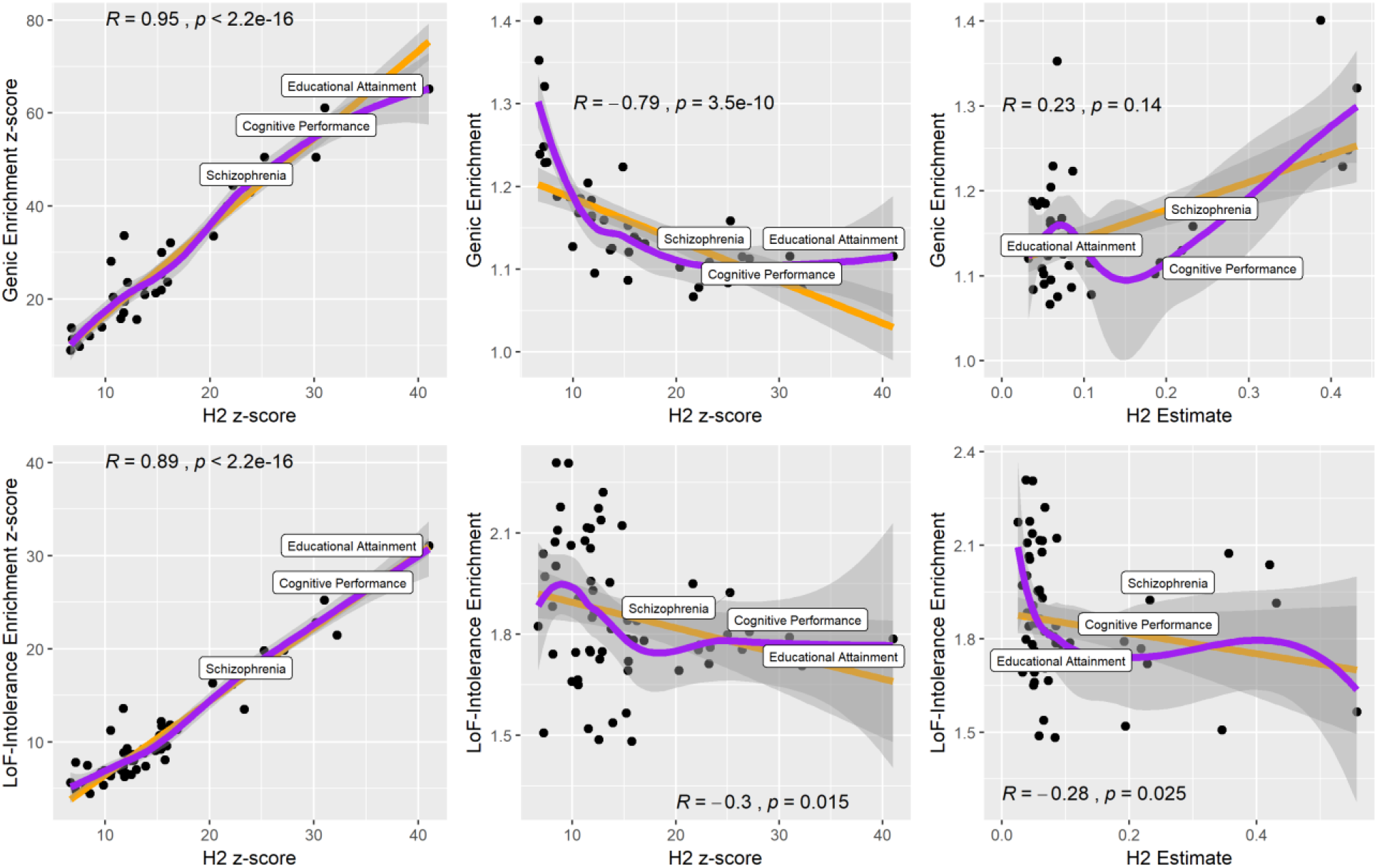
Robust linear (orange) and local (LOESS, purple) relationships between heritability (*h*^*2*^ estimate and *z*-score) and enrichment of *h*^*2*^ attributed to genic and loss-of-function (LoF) intolerant SNPs (estimate and *z*-score) for phenotypes demonstrating nominally significant enrichment of these annotations. The *h*^*2*^ of labeled phenotypes was FDR significantly (*q*<0.05, Table 1) enriched for both genic and LoF intolerant SNPs. Note that robust linear regression down-weighs data points by their distance from the regression line resulting in data point distributions visually inconsistent with a linear model (*e.g.*, middle and right panels in the bottom row).

**Figure S2.**
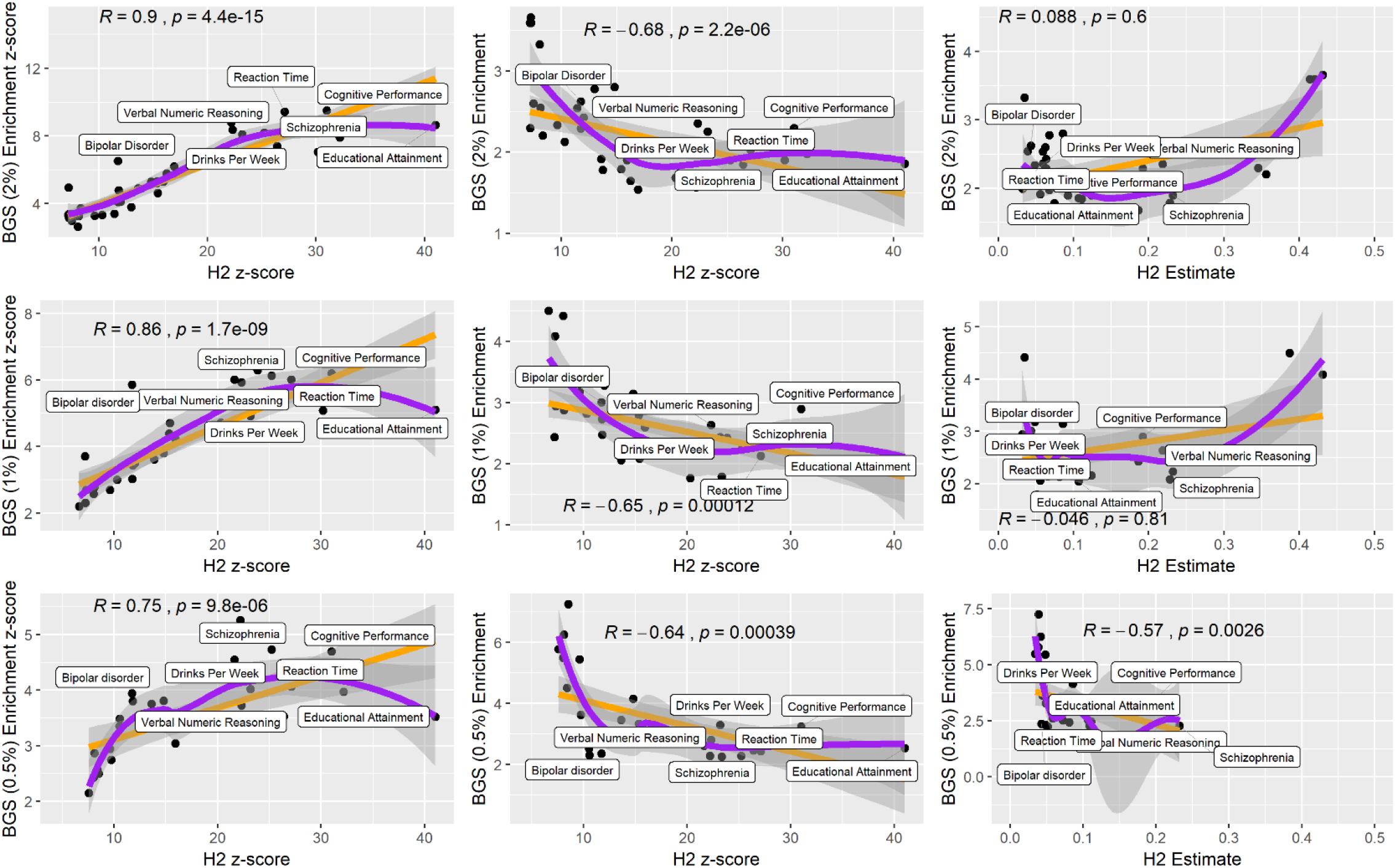
Robust linear (orange) and local (LOESS, purple) relationships between heritability (*h*^*2*^ estimate and *z*-score) and enrichment of *h*^*2*^ attributed to background selection (BGS) intolerant SNPs (estimate and z-score) for phenotypes demonstrating nominally significant enrichment of these annotations. The *h*^*2*^ of labeled phenotypes was FDR significantly (*q*<0.05, Table 2) enriched for both genic and LoF intolerant SNPs.

